# On the role of urban tropical tree collections in carbon allocation: expanding their functions beyond cultural and biodiversity conservation

**DOI:** 10.1101/2024.01.28.577605

**Authors:** Edwin Echeverri-Salazar, Juan Camilo Villegas, Lía Isabel Alviar-Ramírez, Edwin Andrés Mora

## Abstract

Trees support key processes in both natural and managed ecosystems. In highly intervened urban environments, trees have been generally associated with benefits such as air quality, microclimate regulation, and biodiversity conservation. University campuses contain diverse and well-managed tree collections that provide local functions such as education, conservation, research, and landscaping. However, little has been discussed about these collections in the general urban setting and how they relate to other urban ecosystem processes, such as carbon cycling. This is particularly evident in tropical regions where no current urban forest carbon sequestration estimations are available. In this work, we present the results of a pilot estimation of the carbon storage function of the university tree collection at the Universidad de Antioquia (Medellín, Colombia) through a bounding calculation that combines tree inventory data and allometric equations. Our results show that, on average, the university tree collection (including palms) sequesters 3.4 Mg C/ha/year (4.2x10^−2^ Mg C/tree/year). Remarkably, our results are comparable to natural tropical forests, particularly in locations with similar climatic and biophysical conditions. When compared to other urban studies, our estimation ranges between 1.2-20.8 times larger than cities and other urban areas with similar estimations. We present a novel integrative method for estimating carbon storage and sequestration that can be widely applied in information-limited tropical contexts. We discuss how management practices of these urban forests contribute to improving their capacity to store carbon more efficiently and effectively participate in other urban ecosystem processes that derive benefits to society. More generally, our results highlight the role of universities and other similar urban tree collections (i.e. botanical gardens and urban parks) in local and regional ecosystem functions and their potential contribution to global carbon cycling.

## 1. Introduction

Natural forests contain approximately 80% of the continental biodiversity and are key to the functioning of the biosphere, while only covering approximately a third of the world’s continental area (Aerts & Honnay, 2011). However, other types of ecosystems, such as planted forests and particularly urban forests have been also recognized for their ecological value that extends beyond ornamental and aesthetic functions (Carreiro, 2008; Niemelä, 1999; Sukopp, 2002). Extensive research illustrates the growing interest on the social, economical and environmental benefits of urban tree collections (Clark & Matheny, 2009; Dobbs *et al*., 2014; Sarajevs, 2011), including benefits related to their nature and function, such as environmental cooling via reductions in urban heat island effect (Rosenzweig *et al*., 2009), partial and psychological reduction of noise pollution (Clark & Matheny, 2009), UV filtering (Grant *et al*., 2002), reduction of atmospheric pollution and carbon storage (Nowak *et al*., 2006), among others documented in the literature.

The carbon storage function is particularly relevant in the context of global change, particularly in relation to atmospheric concentrations of greenhouse gases and their relation to current and future climate (Solomon *et al*., 2009). Several national and international programs have directed their efforts to promote urban forestation, as a mechanism to reduce air pollution in general, as well as to contribute to greenhouse gas assimilation (Jo & Mcpherson, 1995; Pincetl *et al*., 2013). Multiple studies have assessed the potential for carbon storage in these ecosystems (Mcpherson *et al*. 2006; Nowak *et al*., 2006). Managed urban forests, especially in highly productive areas such as the tropics, can represent an important focus for carbon assimilation and storage. Yet, lacking are quantifications of carbon sequestration potential in tropical urban forests, potentially associated with limitation on monitoring and observation of tree growth in these areas.

In this communication, we present a pilot estimation of the potential carbon assimilation of an urban forest to the tree collection of the University of Antioquia – (UdeA) Medellín, Colombia. We propose a methodology that uses a combination of empirical measurements with theoretical growth relationships to estimate tree growth and carbon sequestration. We contrast our results with available studies in both natural tropical forests and other urban forests (including university tree collections) from different latitudinal and altitudinal locations (mostly in temperate and subtropical areas). We discuss how the diversity of species, ages and the spatial layout of the collection, in combination with efficient maintenance could explain its high potential for carbon sequestration. In addition, we highlight the importance of preserving and expanding these urban forests and to expand their context (educational and ornamental) to additional regulation benefits derived from their function.

## 2. Methods

### 2.1 Location

Our study was developed in the campus of the University of Antioquia, in Medellín – Colombia (6° 16′ 2.7″ N, 75° 34′ 6.2″). The city is located in a narrow intermountain valley (Aburrá valley) at approximately 1560 meters above sea level, mean annual temperature is 24ºC and mean annual precipitation is 1571 mm (Alcaldía-de-Medellín, 2006). The city’s population density is approximately 2556 inhabitants/km^2^, mostly concentrated in the downtown area, where the university campus is located. The 23.75-hectare University campus is recognized locally as an important biodiversity hotspot, as well as heat island mitigation area (Appendix Fig. 1).

### 2.2 Tree inventory

We used the 2010 university tree and shrub inventory (Vélez, 2010– internal University facilities management document) that includes a total of 2282 individuals. Among those, DBH (diameter at breast height – tree diameter measured at 1.3 meters above the surface) is reported for 1791 trees; and stem-height for 157 palms instead of DBH. When trees had multiple trunks, all trunks ≥10 cm were measured and considered an individual trunk; DBH peaked around 25 cm, with values ranging between 10 cm and > 200 cm (Fig 1A) for the larger trees (*Ficus benjamina*). In our study, we included all 1948 individuals with DBH≥10.

**Figure 1.**
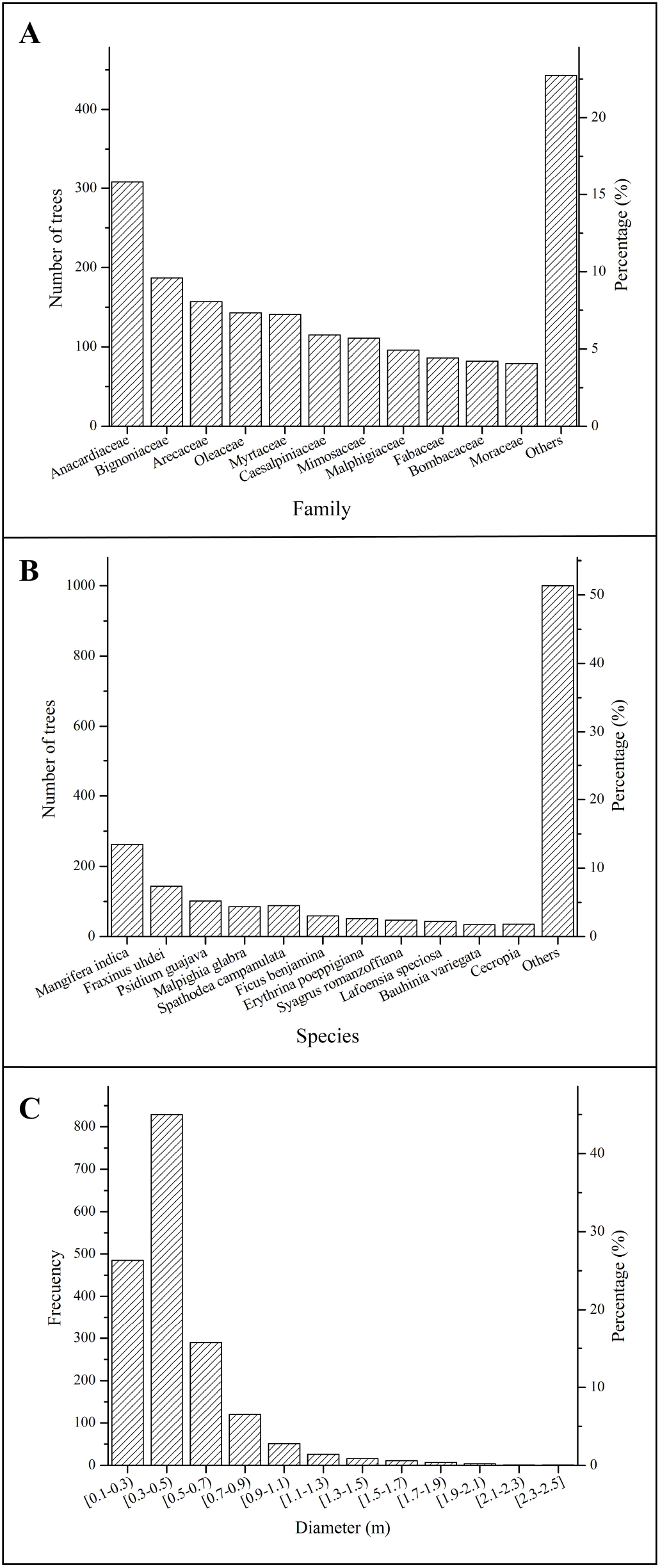
General biometric and taxonomical characteristics of the University tree collection. (A) DBH (diameter at breast height) size distribution. (B) Common families and (C) Species in the university tree collection.

Our sample was composed mostly by angiosperms (1940 individuals in 41 families) and gymnosperms (8 individuals in two families). The most common families in the collection include: Anacardiaceae, Bignoniaceae, Arecaceae (among others presented in Fig. 1B), with the most common species being *Mangifera indica* (mango tree – non-native), *Fraxinus uhdei*, and *Psidium guajava* (guava tree), with over 100 individuals each (these and other common species are presented in Fig 1C). Most of these common species are common flowering and fruiting species that support a highly diverse array of fauna within the University campus.

### 2.3 Biomass and carbon sequestration estimations

The amount of carbon stored in the tree collection was estimated with the allometric equations proposed by Sierra *et al*. (2007). These equations have been validated for secondary forests in Colombia, with climatic and taxonomic distributions comparable to those found in the campus tree collection. The equations use DBH (in cm) to estimate aboveground biomass (AB; in kg – Equation 1) and belowground biomass (BGB; in kg – Equation 2). We report both components as well as an estimated wet Total Biomass (TB; in kg – Equation 3). Given the effects of management and site characteristic (particularly soil conditions) being different from natural forests, we did not consider fine root biomass or vines due to uncertainty in this estimation.

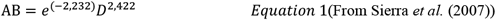

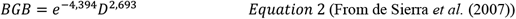

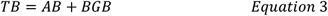

Palm tree biomass (PB; in kg – Equation 4) was also estimated from allometric equations proposed by Sierra *et al*. (2007), for aboveground biomass as a function of trunk height (H -in m) (Eq. 4)

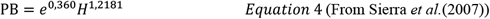

To quantify carbon sequestration, it is necessary to estimate growth for a period of time. To do so, we used the inventory data described above in conjunction with scaling allometric relations proposed by Enquist *et al*. (2007). These equations estimate biomass change in time as follows (Eq. 5)

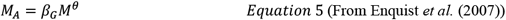

Where *M*_*A*_ is the annual production of biomass (g year^-1^); *β*_*G*_ is an allometric constant corresponding to the net biomass production per unit foliar mass, and its values vary for angiosperms (2.43g^1/4^year^-1^,IC_95%_=0.44-11.92) and gymnosperms (1.35g^1/4^year^-1^,IC_95%_=0.41-4.42), originally developed through integration of global plant trait databases (Enquist *et al*., 2007 – Supplementary material); *M* corresponds to total tree biomass (from equations 3 and 4); and *θ* is a metabolic scaling parameter, associated with the geometry and structural characteristics of trees, for which a value of 3/4 is assigned, according to metabolic scaling theory that provides a metabolic framework that can be generally applied to all tree forms (Enquist *et al*., 2007). To convert biomass into carbon units, we use the factor of 0.42 for palms (Vlek *et al*.,2005) and 0.45 to the other trees (Sierra *et al*., 2007). A conceptual model of our overall methodology is presented in Appendix Fig 2.

To compare and contextualize our estimations, we searched in the existing literature carbon sequestration estimations for three types of conditions that included: (i) studies in tropical forests in general, (ii) studies in tropical areas with similar climate and altitudinal conditions, and (iii) other studies in urban settings, including cities and university campuses.

## 3. Results and Discussion

Total biomass in the University collection for 2010 tree inventory data was 3,751.66 tons of Carbon (corresponding to 13,476.80 Mg CO_2_) with a sequestration of 80.87 MgC/year (294.89 Mg CO_2_/year) (CI_95_=14.66-396.90) (Table 1) and sequestration per unit of area of 3.41 Mg C ha^-1^year^-1^ (12.51 Mg CO_2_ ha^-1^year^-1^). Most of the sequestration (89.4%) occurs in trees belonging to the 10 families in the collection (Fig 2A), 8 of which are among the most common (Fig. 1A). More specifically, most of the sequestration occurs in individuals of the *Moreceae* family, which, although not the most common, includes individuals with the largest sizes in the collection (*Ficus benjamina, Ficus elastica* with diameters ranging from 0.34-2.00m and 0.37-3.98 m, respectively), This observation agrees with recent observations that larger/mature trees have greater potential for carbon sequestration than smaller trees (Stephenson *et al*., 2014). Conversely, total carbon sequestration was greater in small and medium size trees (DBH size class 0.1-0.9 m – Fig. 2B, because in these diametric classes can we find 95% of the trees.

**Table 1.**
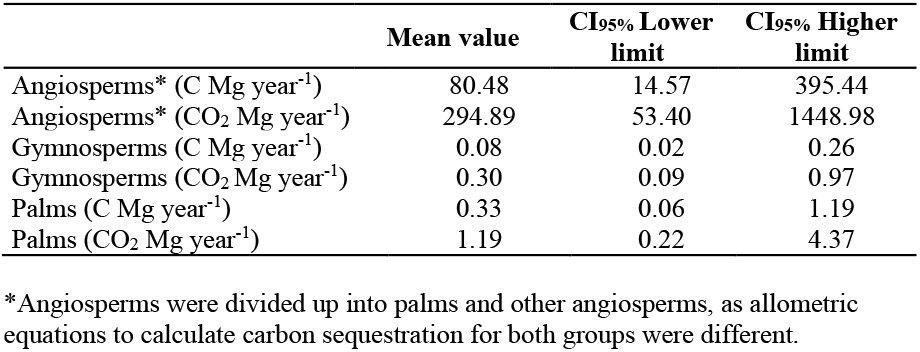
Carbon and CO_2_ sequestration in Angiosperms and Gymnosperms.

|  | Mean value | CI95% Lower limit | CI95% Higher limit |
| --- | --- | --- | --- |
| Angiosperms* (C Mg year<sup>-1</sup>) | 80.48 | 14.57 | 395.44 |
| Angiosperms* (CO<sub>2</sub> Mg year<sup>-1</sup>) | 294.89 | 53.40 | 1448.98 |
| Gymnosperms (C Mg year<sup>-1</sup>) | 0.08 | 0.02 | 0.26 |
| Gymnosperms (CO<sub>2</sub> Mg year<sup>-1</sup>) | 0.30 | 0.09 | 0.97 |
| Palms (C Mg year<sup>-1</sup>) | 0.33 | 0.06 | 1.19 |
| Palms (CO<sub>2</sub> Mg year<sup>-1</sup>) | 1.19 | 0.22 | 4.37 |

\*Angiosperms were divided up into palms and other angiosperms, as allometric equations to calculate carbon sequestration for both groups were different.

**Figure 2.**
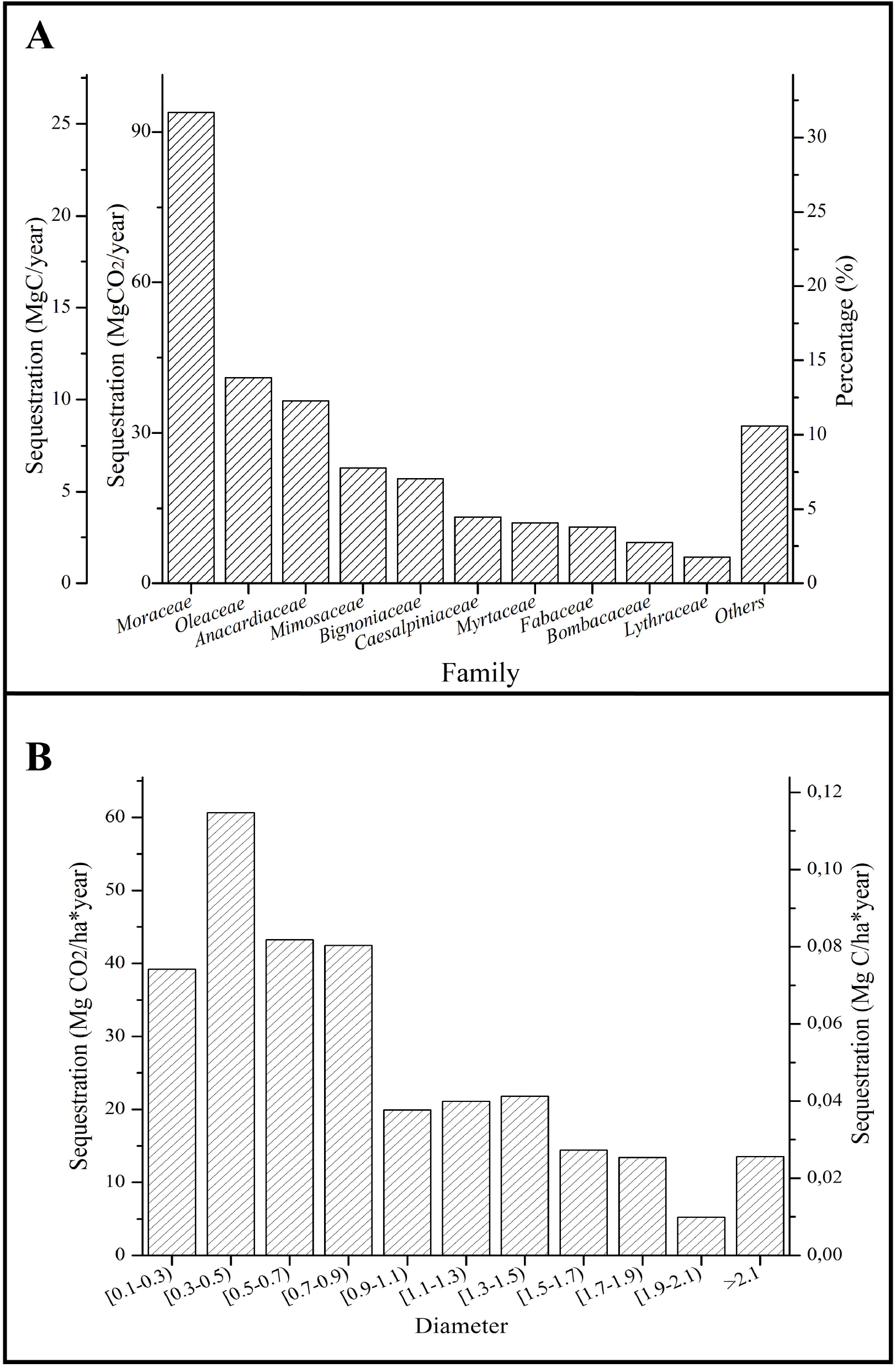
Carbon sequestration per family per year (left axis) and as a percentage from the total sequestration (80.87 Mg C per year) in the collection (right axis), including the ten families with the largest amounts of carbon sequestration.

Our estimations are not predictive in nature, as there is uncertainty in both tree biometric measurements (i.e. not including buttress measurements in our estimations which could lead to underestimations of as much as 10% in DBH (Metcalf *et al*., 2009), as well as uncertainty in biomass and tree growth calculations from allometric equations. For instance, we have used a single equation for palm tree growth, where other studies have shown that growth in this group can be species-specific (Goodman *et al*., 2013). Similarly, by using general equations in both the estimation of carbon storage and sequestration, we may be overgeneralizing species (or family) specific ecological traits. However, we selected the most relevant allometric models to our study site, allowing us to limit uncertainty in the calculations. Our proposed method is useful to overcome data limitations in many environments (particularly the tropcis) and provide an illustration of the necessity to incorporate urban forests into the global carbon dialogue.

In general, our results are comparable to other studies in natural tropical forests that have found similar amounts of carbon sequestration in tropical regions (Steininger, 2000; Worbes & Raschke, 2012, Fig. 3A). Most of these studies calculated aboveground carbon sequestration, still comparable with our estimation of aboveground carbon sequestration, with greater amounts in lowland tropical forests (Bolivia and Brazil) and lower amounts in higher elevation and secondary forests. Notably, when compared to similar published studies in urban forests (Aguaron & Mcpherson, 2012; Liu & Li, 2012; Velasco *et al*., 2013) and university tree collections (Cox, 2012; De Villiers *et al*. 2014) in temperate and subtropical regions of the world (Fig. 3B), our results indicate that our tree collection has the potential to incorporate as much as 1.2 to 20.8 times the amount in other similar urban tree collections. These results highlight the potential of tropical managed tree collections to be an important carbon sink. Carbon sequestration in both natural and urban forests depends directly on tree size and density (amount of trees per unit area); to account for these—and to make our results more comparable to other studies in urban systems—we calculated carbon storage per unit tree. When considered in a per unit tree basis (Fig. 3B left axis), our estimations indicate that our tropical tree collection can store from 2.5 to 56.7 times more carbon than similar urban forests.

**Figure 3.**
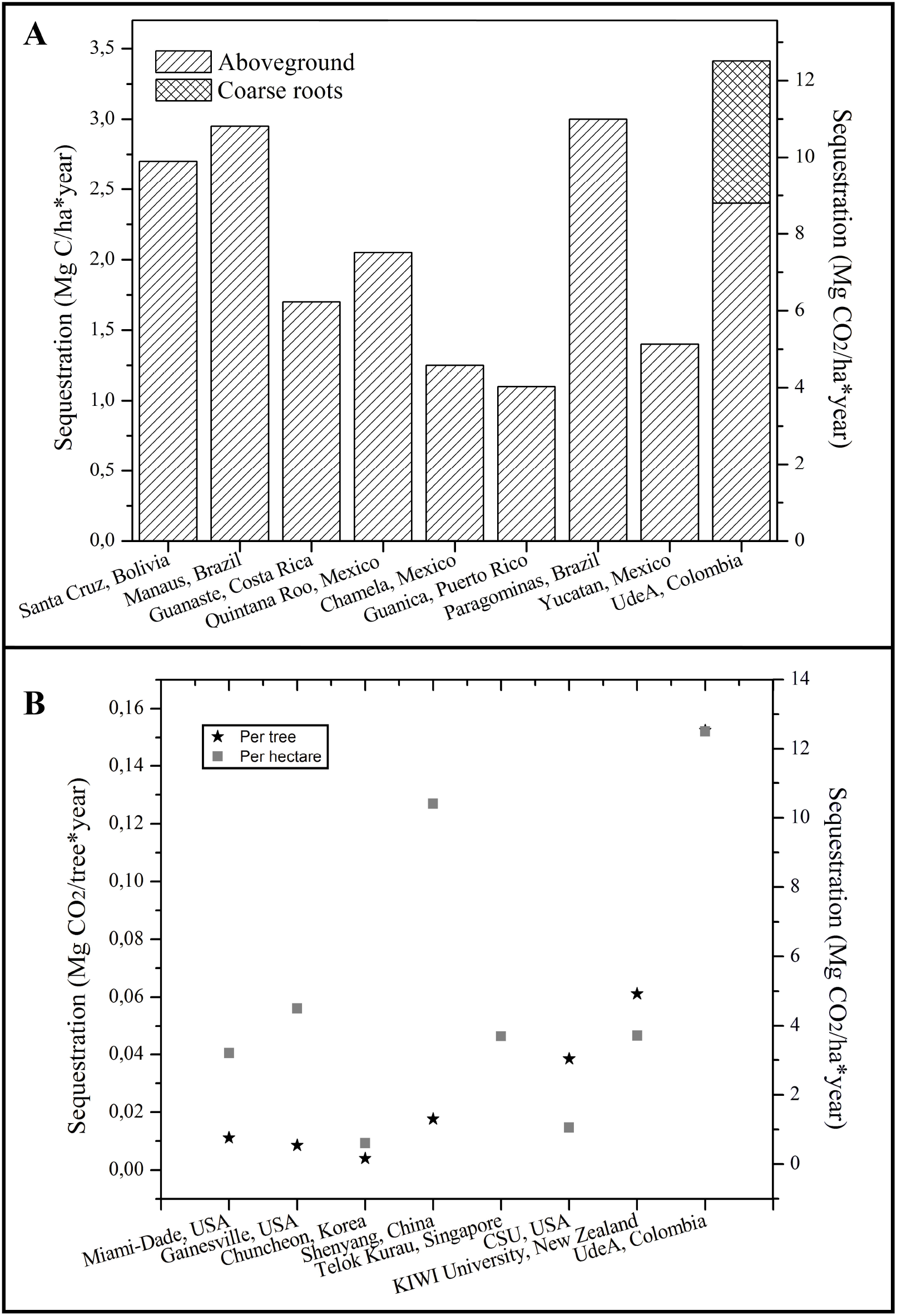
Carbon sequestration in the University tree collection in comparison with (A) tropical forests (data from Steininger, 2000; Worbes & Raschke, 2012) and (B) other urban forests, including city and University tree collections (data from Aguaron & Mcpherson, 2012; Liu & Li, 2012; Velasco *et al*., 2013; Cox, 2012; De Villiers *et al*. 2014)

The University campus is located in a highly urbanized area with high potential for air pollution associated with vehicle emissions, and recognized as one of the most critical areas in CO concentrations in the city (Daniels *et al*., 2007). Although not considered in our study, other studies have shown that the high concentration of gases, in densely populated urban areas has the potential to enhance tree growth (Gregg *et al*., 2003; Keeling *et al*.,1996). However, it is widely recognized urban trees may not be able to offset all emissions. In general, efforts to control gas emissions in cities should be devoted to decrease emission, while increasing the potential for mitigation through urban forestry (Brack, 2002).

Our estimations of carbon sequestration in the University tree collection are among the first attempts to estimate the potential role of tropical urban forests in carbon sequestration. Our results are an invitation to the research community to more explicitly consider the potential carbon benefits of tropical urban forests, along with the already recognized list of benefits from urban trees (Clark & Matheny, 2009; Sarajevs, 2011). Notably, we propose a novel approach based on existing data and allometric equations that can be generally used to provide estimations of carbon storage and sequestration in urban environments with limited available data.

Collectively, our results—which are intended to illustrate the potential of urban tropical tree collections for storing carbon—illustrate that the amount of carbon sequestration in tropical urban tree collections is comparable to other tropical environments, which in turn are significantly higher than estimations for other urban ecosystems in temperate regions. We hypothesize that these potential is associated with higher availability of solar radiation throughout the year (with no marked seasonality) and effective maintenance that compensates for potential nutrient limitations with fertilization (in our case made with in-situ composted material) as well as irrigation systems compensating for potential water limitations during the drier portions of the year. More generally, our results highlight the role of urban tree collections (i.e. university campuses, botanical gardens and urban parks) in local and regional, and potentially, global carbon cycling, along with their already recognized function in educational, cultural and biodiversity support services. We encourage the research community to collectively improve our ability to better characterize tropical urban forest function and to incorporate it into the global carbon management coordination as suggested by the recent 21^st^ conference of the parties (COP21).

## Acknowledgements

We thank Gladys Vélez for tree biometry data, Dirección de Gestión logísitca e infraestructura Universidad de Antioquia for logistical support, Grupo Aliados con el Planeta for conceptual discussion, Katerine Cárdenas, Ana Posada, Alejandro Montealegre, Andrés González, Jorge Giraldo and Kevin Pulgarín for field support. Financial support for this work was provided by Programa jóvenes investigadores Universidad de Antioquia 2012-2013, GIGA research group, and Estrategia de sostenibilidad 2014-2015 Universidad de Antioquia.

---

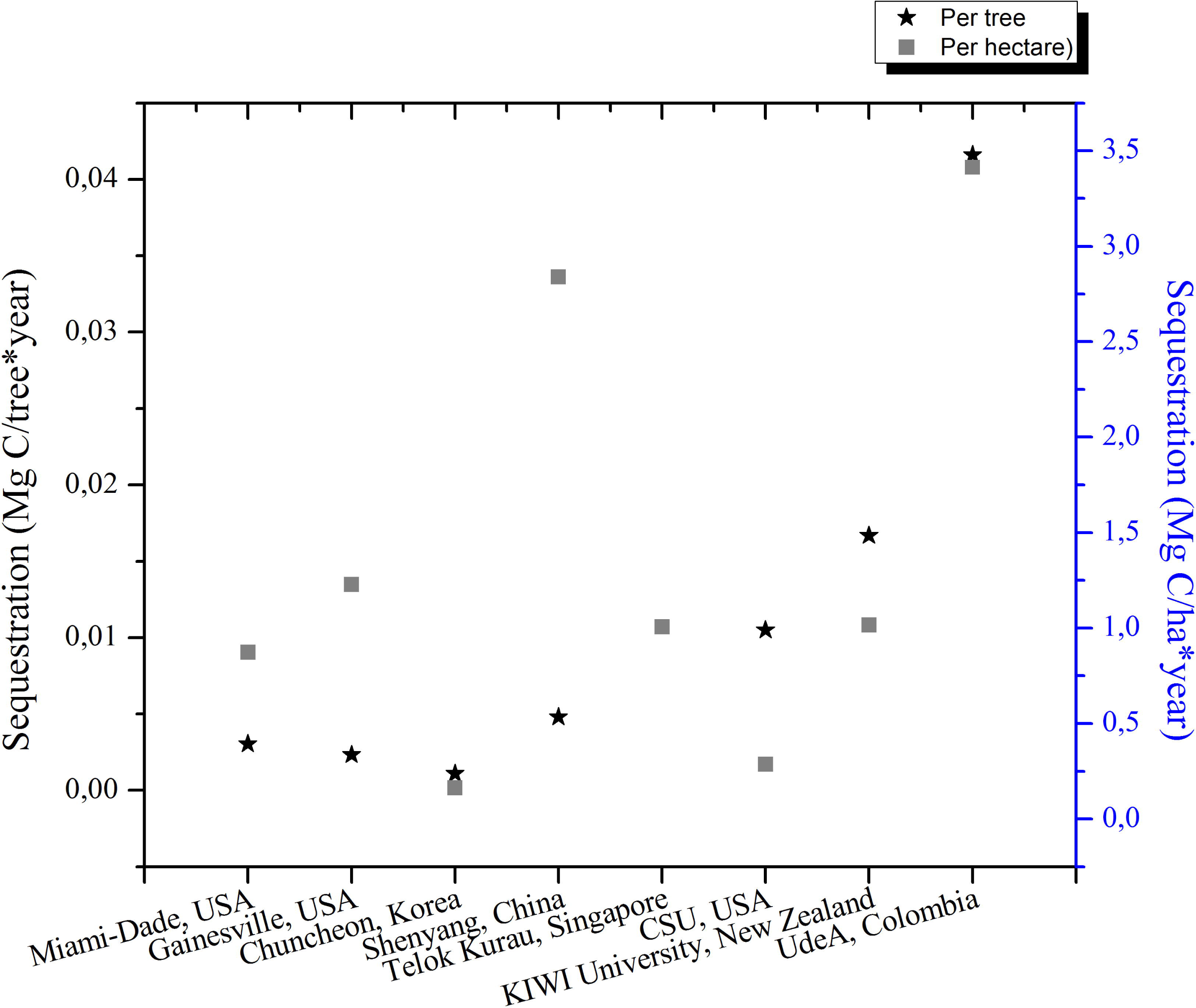

