## Appendix Figures for "On the role of urban tropical tree collections in carbon allocation: expanding their functions beyond cultural and biodiversity conservation"

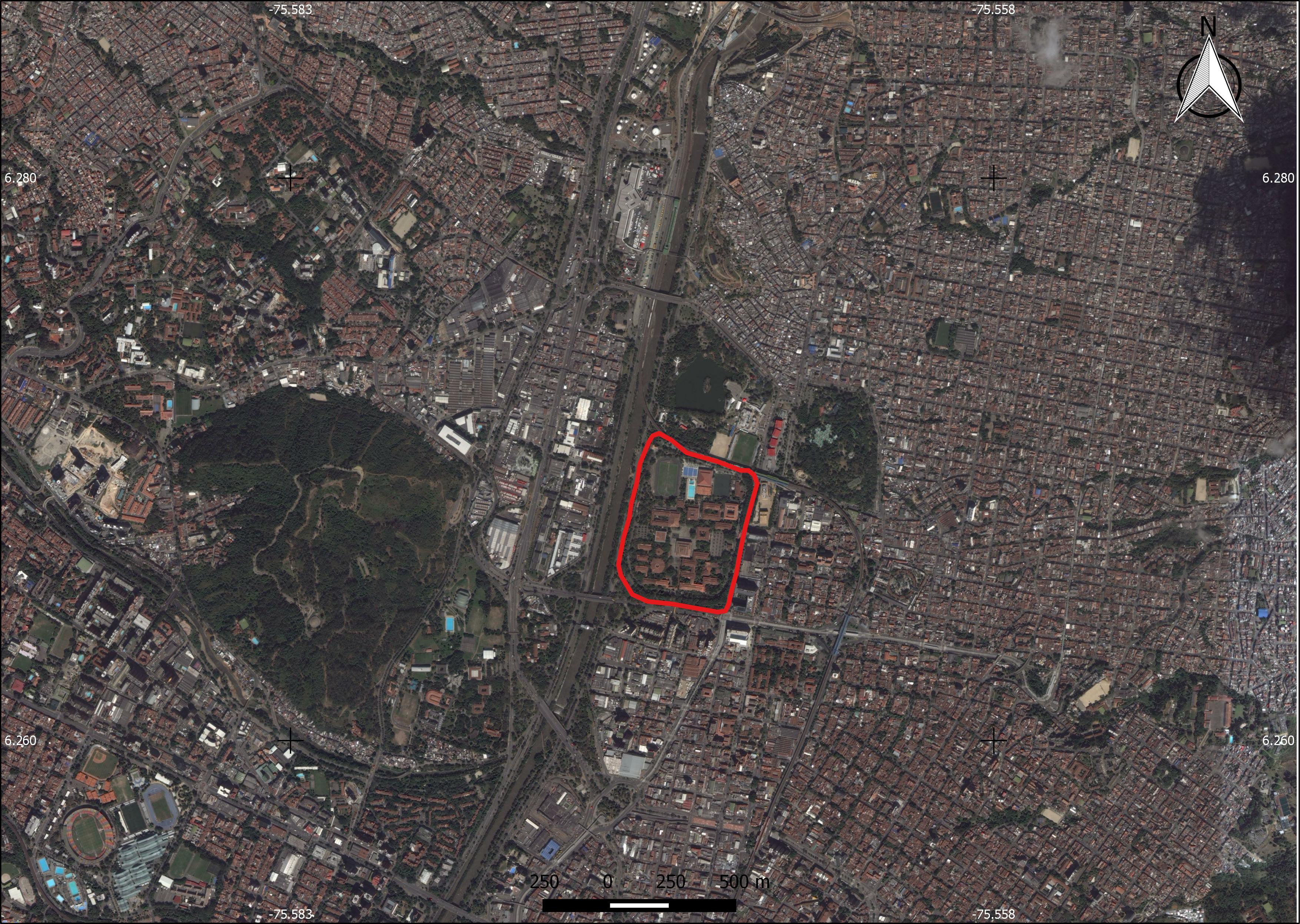

Appendix Figure 1. General Location of the Universidad de Antioquia campus in Medellín, Colombia. Image from Google Earth

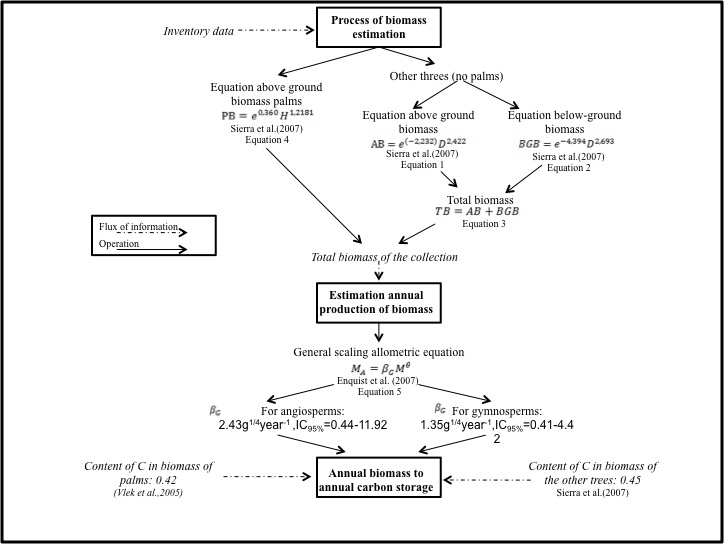

Appendix Figure 2- General Methodological approach for the estimation of carbon sequestration in tropical urban forests – case study Universidad de Antioquia
